# TE-Seq: A Transposable Element Annotation and RNA-Seq Pipeline

**DOI:** 10.1101/2024.10.11.617912

**Authors:** Maxfield M.G. Kelsey, Radha L. Kalekar, John M. Sedivy

## Abstract

The recognition that transposable elements (TEs) play important roles in many biological processes has elicited growing interest in analyzing sequencing data derived from this mobile ‘dark genome’. The TE-Seq pipeline conducts an end-to-end analysis of RNA-sequencing data, examining both genes and TEs. It implements the most current computational methods tailor-made for TEs, enabling a comprehensive analysis of TE expression at both the individual element level and at the TE clade level. If supplied with long-read DNA sequencing data, it creates a TE-complete genome incorporating non-reference (polymorphic) TE-loci, enabling the functional characterization of the evolutionarily youngest mobile elements in the genome.

## Background

Transposable elements (TEs) are genetic elements with the unique ability to move throughout genomes. These processes can increase TE genomic copy number, so much so that TEs comprise roughly half of the human genome and similarly large fractions of other mammalian and eukaryotic genomes [1]. TEs are important agents of genome evolution by promoting a variety of genetic events, such as gene inactivation, complex DNA rearrangements, mobilization of regulatory sequences, and others [2, 3]. TE-mediated evolution remains operant in all species examined including humans [4, 5]. Through adaptive evolution, TE-derived sequences have become involved in many fundamental cellular processes, such as chromosome segregation, gene regulation, and telomere function [1, 6]. Retrotransposable elements (RTEs), in particular Long Interspersed Nuclear Elements (LINEs), Short Interspersed Nuclear Elements (SINEs) and Endogenous Retroviruses (ERVs) are of considerable interest due to their abundance and activity in many mammalian genomes.

Until recently, RTE activity in somatic tissues has received little attention. While these events are not inherited from generation to generation, they can exert deleterious effects on the cells and tissues in which they occur, such as the promotion of cancer, inflammation and neurodegeneration [7–10]. One important mechanism is genome instability and DNA damage caused by the retrotransposition process [11]. A more recently appreciated mechanism is the stimulation of innate immunity antiviral defenses by RTE nucleic acids [8].

While the broad relevance of TEs is clear, the repetitive nature of their sequences imposes analytical challenges, and hence they are often neglected in RNA/DNA-Seq workflows and analyses [12]. This pipeline (Fig. 1) aims to render TE investigation more tractable by addressing concerns pertaining to: i) imperfect alignment of repetitive elements, ii) non-reference elements, iii) non-autonomous transcription of TEs driven by adjacent genes, and iv) the quality of TE annotations.

**Fig. 1.**
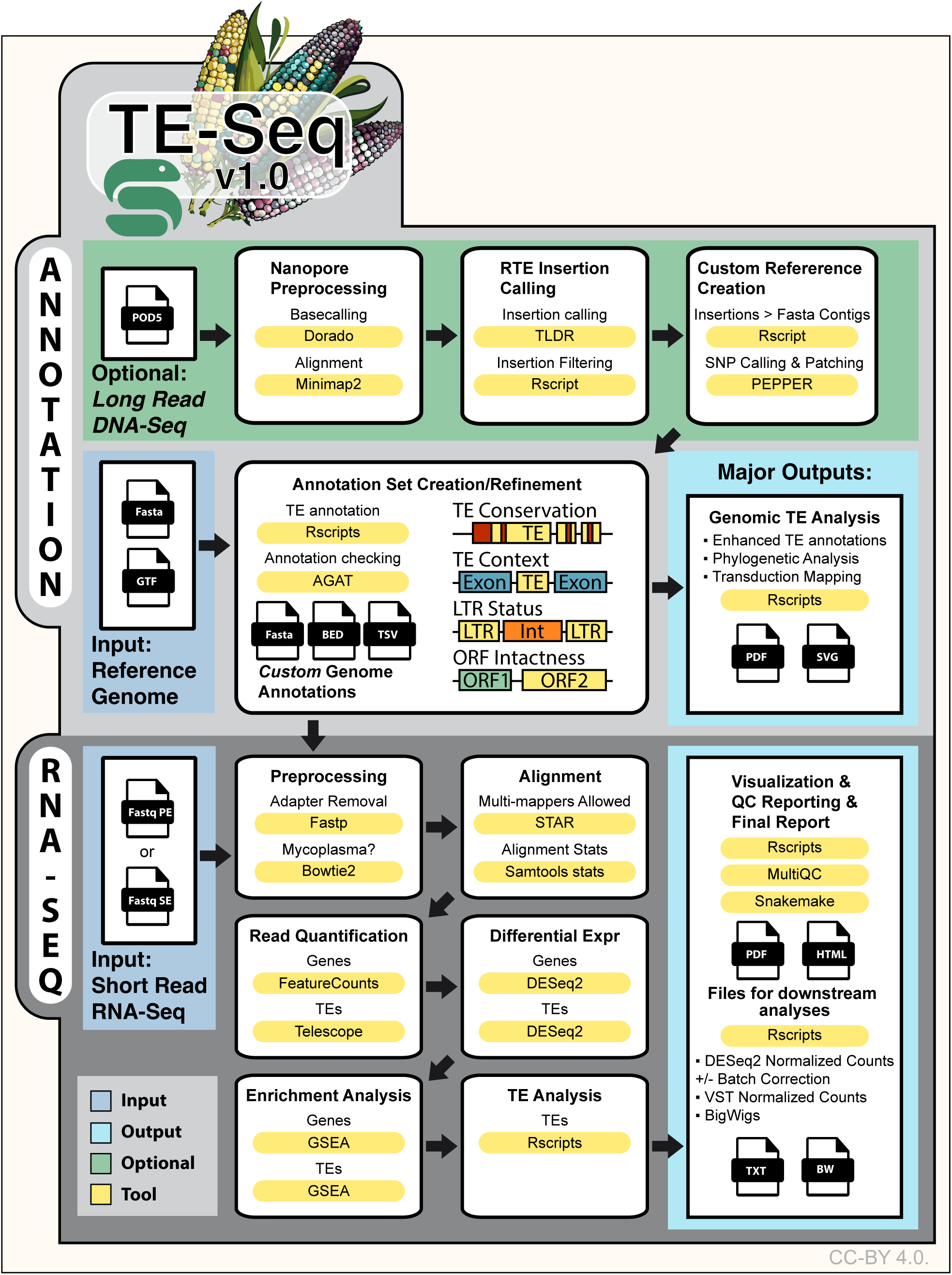
TE-Seq pipeline overview.

RTEs are classified into superfamilies (e.g. LINE), families (e.g. LINE-1), and subfamilies (e.g. L1HS, specific for *Homo sapiens*). RTE count quantification has to confront the issue of multimapping reads: if two insertions in a genome are identical in sequence, it is impossible to uniquely assign reads derived from them uniquely to either of locus. In practice, however, almost all RTE copies contain some polymorphic sites (such as SNPs or indels), allowing some fraction of their reads to be uniquely mappable. A strategy of examining only uniquely mapped reads may at first glance appear attractive, but this approach is biased against evolutionarily young RTE elements, which are less polymorphic than older elements [13, 14]. However, young RTE subfamilies are of greatest interest because they are the most mobile, and thus tools have been developed to handle reads derived from non-uniquely mappable sequences.

This problem was recognized early on and programs such as TEtools randomly assigned a multi-mapper read to one of its mapped loci; this greatly improved quantification accuracy [15]. Other tools such as RepEnrich [16] focused on providing accurate assessments of expression at the TE subfamily level by pooling all uniquely mapped reads to a given TE subfamily, and remapping multi-mappers to TE subfamily consensus sequences. As sequencing read lengths improved, with 150 bp paired-end reads becoming standard, the yield of uniquely mapping reads increased, and new strategies to deal with the multi-mapping problem were developed, such as TElocal (https://github.com/mhammell-laboratory/TElocal), L1EM [17], and Telescope [18]. These programs implemented iterative, expectation maximization-based assignments of multi-mappers to single loci. They rely on the assumption that loci which have uniquely mapped reads are likely truly expressed, and therefore multi-mapping reads are iteratively assigned to TE loci which accrue the largest number of uniquely mapped reads. Our TE-Seq pipeline (Fig. 1) utilizes the Telescope tool since it accepts user-provided gene-transfer format (GTF) formatted TE annotations, making it more flexible than either TElocal or L1EM; the former relies on pre-built custom-TE indices, while the latter focuses solely on LINE-1 elements.

A complete account of every TE insertion in a genome would greatly enhance the quality and interpretability of TE RNA-Seq results. Currently mobile RTE subfamilies, such as L1HS, are constantly evolving and generating new germline insertions. Each human individual has approximately 100 L1 insertions that are not captured by a given reference genome (e.g. Hg38). As a result, reads derived from these elements will inevitably be misassigned to other reference elements. This problem is particularly acute since the polymorphic, non-reference RTEs are likely some of the most active elements in the genome. As a solution to this problem, our TE-Seq pipeline allows for the optional injection of Nanopore DNA sequencing data in order to call non-reference TE insertions and include them in downstream analyses (Fig. 1).

Recognition of an element’s ‘genic context’, or its spatial relationship to cellular genes, helps to estimate potential regulatory influences between TEs and the surrounding host genome, in particular, whether an element was autonomously expressed (transcribed by its own promoter) or whether it was passively transcribed by gene read-through. The latter possibility is diminished if no adjacent genes exist. To this end, the TE-Seq pipeline categorizes elements according to their relationship with cellular genes (Fig. 1). This information serves another diagnostic function: if libraries differ greatly in terms of intronic content (a potential technical artifact owing to differences in sample preparation), this can lead to a spurious apparent increase in TE expression. The degree to which this pitfall may affect a dataset is assessed by comparing changes in intergenic-TE expression to changes in intronic-TE expression.

## Results

In order to catalogue all non-reference RTE insertions in LF1 human primary lung fibroblast cells [19], we performed Nanopore DNA sequencing. Sequencing coverage and N50 values were high (>60X, >24 kb). We called non-reference insertions using the TLDR program [20], and retained only those insertions passing both TLDR and our in-house quality filters (Methods). We observed 226 Alu insertions, 117 L1 insertions, and 24 SVA insertions (Fig. 2A). While some L1 insertions were truncated, we found 65 full-length L1HS insertions, of which 46 were intact with conserved ORFs. Most insertions were intergenic or intronic (Fig. 2B). A smaller fraction were either adjacent to coding or non-coding genes, or found within an exon or a non-coding transcript. Compared to reference L1HS elements, non-reference L1HS elements were much more likely to be intact, and non-reference elements made up 30% (46/151) of all intact L1HS elements (Fig 2C). When non-reference intact L1HS consensus sequences were phylogenetically compared to reference intact L1HS elements, non-reference insertions were dispersed throughout the phylogeny (Fig. 2D). Assuming that the most phylogenetically proximal reference element is the most likely non-reference progenitor, a small number of reference intact L1HS elements were predicted to account for the majority of non-reference insertions (Fig. 2E). Under this assumption, the L1HS element (17_q24.2_2) produced eight novel insertions, indicating it may be a particularly active element in the germline. We also generated a pie chart showing the fraction of this genome occupied by repetitive elements (Fig. 2F).

**Fig. 2.**
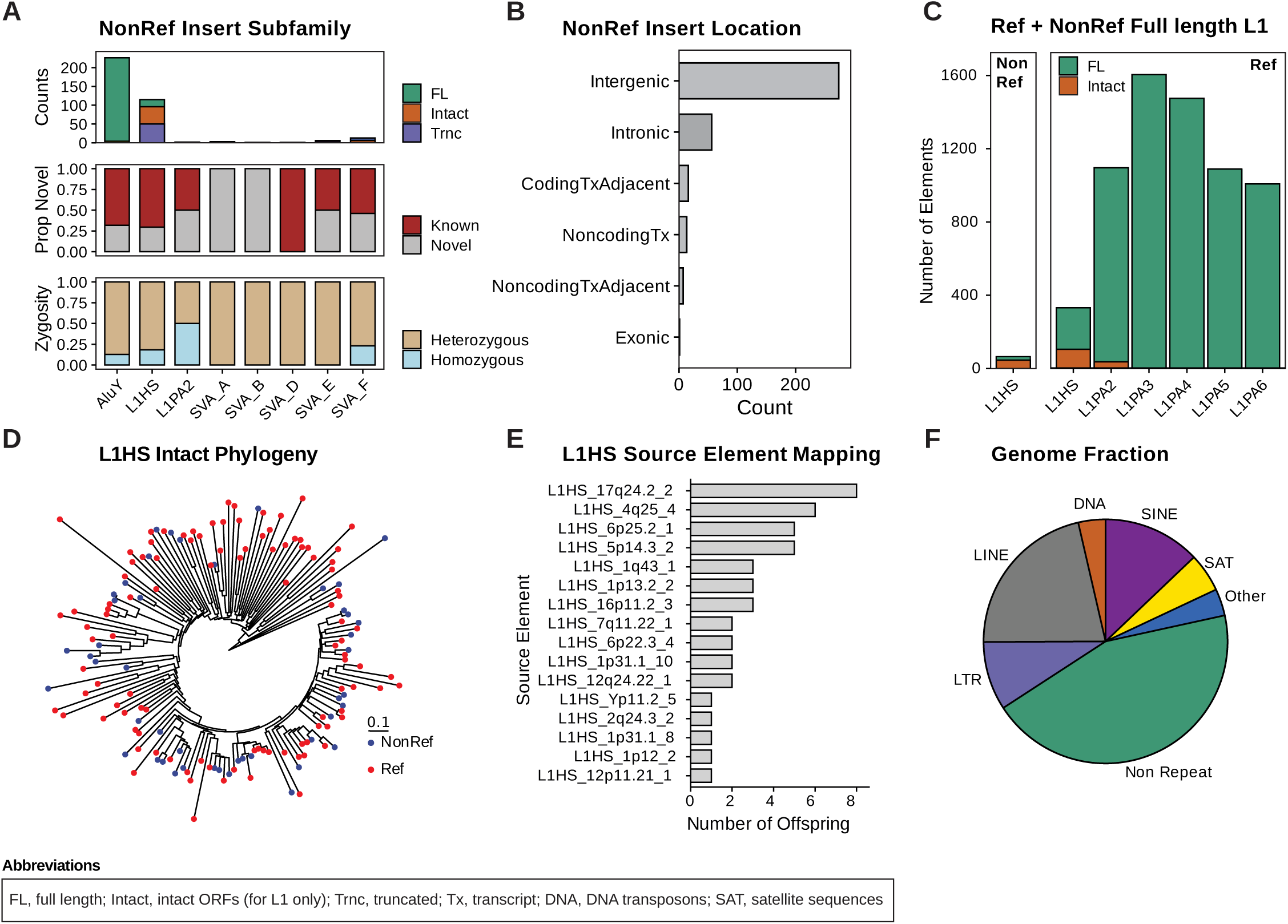
Nanopore DNA-Seq characterization of non-reference RTE insertions in a human genome. TE insertions in LF1 cells were called relative to the telomere-to-telomere T2T-HS1 human reference genome using TLDR. **A** Insertion counts for Alu, L1 and SVA subfamilies (no non-reference ERVs were found). Insertions denoted as ‘known’ have been previously identified in human populations. **B** Insertion positional relationships to cellular genes (for all non-reference RTEs in panel (A)). 5’ and 3’ UTRs are included in the exonic category. **C** Reference versus non-reference full-length L1HS and L1PA2 elements. **D** Phylogeny of intact L1HS elements. Tips are colored according to reference status. **E** Source elements were inferred for all non-reference L1HS insertions on the basis of phylogenetic relationships, and the number of offspring was tabulated for each source element. **F** Genomic occupancy for major clades of repetitive sequences.

Starting with the telomere-to-telomere human reference genome, T2T-HS1 [21], we then created a custom LF1 genome that incorporates quality-filtered TE insertions, rendering them alignable and hence open to scrutiny by RNA-Seq analysis. Enhanced TE annotations were then created to facilitate a thorough investigation of these elements. We applied our pipeline to RNA-Seq data derived from LF1 human primary lung fibroblast cells [19] undergoing senescence. LF1 cells were passaged to replicative senescence, and were harvested 4 and 16 weeks later, yielding early senescence (ESEN) and late senescence (LSEN) conditions, respectively. Early passage proliferating LF1 cells (PRO) were used as a control. Poly-A selected, stranded, 150bp paired-end libraries were sequenced in quadruplicate.

Principal component analysis revealed samples mostly clustered by biological condition (Fig. 3A), excepting one ESEN sample, which clustered closer to LSEN samples. Senescence status neatly divided the first principal component, which accounts for 72.63% of total variation. We observed 7341/6245 up/down-regulated genes in ESEN and 7274/6117 up/down-regulated genes in LSEN (Fig. 3B). While most differentially expressed genes were shared between both senescent conditions, roughly 40% of DEGs were unique to one condition, suggesting biological differences between early and late senescence. Hallmark gene set enrichments by GSEA (Fig. 3C) were consistent with previous reports demonstrating heightened inflammatory signatures and P53 pathway activity, as well as diminished activity of pathways associated with proliferation, including E2F targets, and G2M checkpoint [22]. Canonical transcriptional markers of senescence were significantly elevated in ESEN and LSEN, including CCL2, CDKN1A (P21), CDKN2A (P16), and IL6 (Fig. 3D). The SenMayo gene set was enriched in both ESEN and LSEN (Fig. 3E). Interestingly, hierarchical clustering of SenMayo gene expression showed that this signature was dynamic in senescence, with some SenMayo genes being differentially expressed between ESEN and LSEN (Fig. 3F). The direction of most changes observed in ESEN was preserved in LSEN, but the magnitude of these changes was modulated in LSEN. One large block of genes (top third of heatmap) was highly upregulated in ESEN and comparably modestly upregulated in LSEN. While STRING analysis did not detect functional enrichments in this block compared to the broader SenMayo set, we noted that many of these genes were cytokines (IL1A, IL1B, and IL6) and chemokines (CCL2, CXCL8, CXCL10). Another large block (middle of heatmap) was upregulated modestly in ESEN and much more strongly in LSEN.

**Fig. 3.**
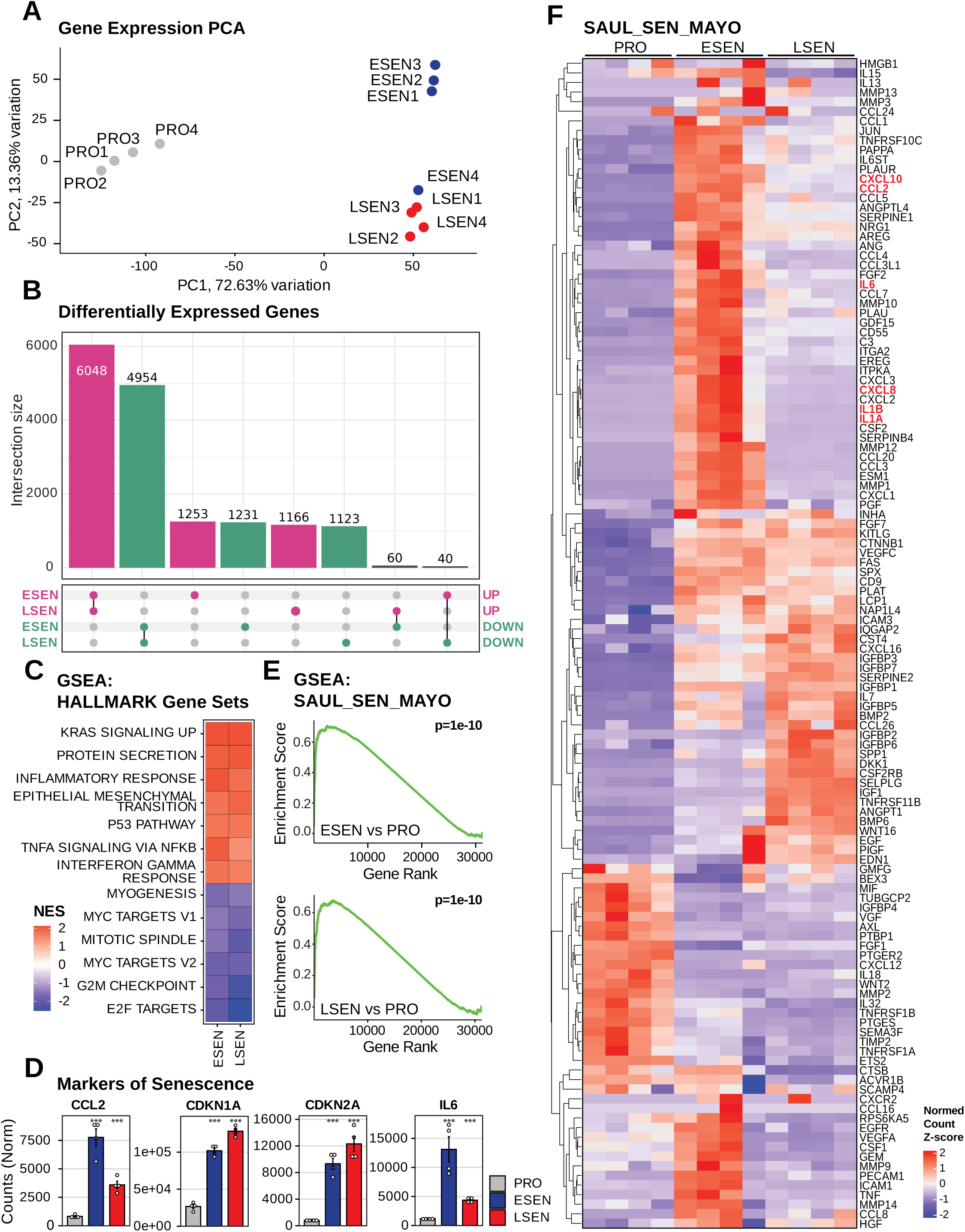
RNA-Seq analysis of cellular genes. Human LF1 cells were passaged until replicative senescence and harvested 4 (ESEN) or 16 (LSEN) weeks later. Early passage proliferating (PRO) LF1 cells were used as control. **A** PCA biplot of cellular genes. **B** Upset plot of differentially expressed genes. **C** Gene set enrichment analysis (GSEA) of MsigDb Hallmark gene set collection for ESEN vs PRO and LSEN vs PRO. **D** Bar plots showing differentially expressed markers of senescence. **E** GSEA plots of the SenMayo gene set for ESEN vs PRO and LSEN vs PRO. **F** Heatmap showing expression Z-scores for all SenMayo gene set members. Genes referred to in the main text are bolded in red. Unless otherwise stated, statistical significance was assessed by means of a two-sided t-test, with panel-level FDR corrected p-values <= 0.05 **\***, <=0.01 **\*\***, and <=0.001 **\*\*\***.

The majority of differentially expressed repetitive elements were upregulated: 10,204 were upregulated in ESEN and 7,418 were upregulated in LSEN, while 4,766 were downregulated in ESEN and 6,206 were downregulated in LSEN. Nearly half of the up- and downregulated TEs were shared between senescent conditions (Fig. 4A). At the superfamily level, RTEs (LINEs, SINEs, LTRs) and DNA transposons were upregulated, while retroposons and satellites were downregulated (Fig. 4B). An examination at the RTE family level found that L1 elements were particularly highly expressed during early and late senescence (Fig. 4C). This observation was further borne out upon examining expression at the RTE subfamily level, where evolutionarily recent L1 subfamilies saw the largest expression fold changes in early and late senescence (Fig. 4D). Gene set enrichment analysis of these RTE subfamilies (Fig 4F), in which subfamilies were treated as gene sets, was concordant with the log2 fold change aggregated count analysis (Fig. 4D). Both showed that evolutionarily recent L1 and HERV-int subfamilies were enriched in ESEN and LSEN, while SVA sets were depleted (Fig. 4D, F).

**Fig. 4.**
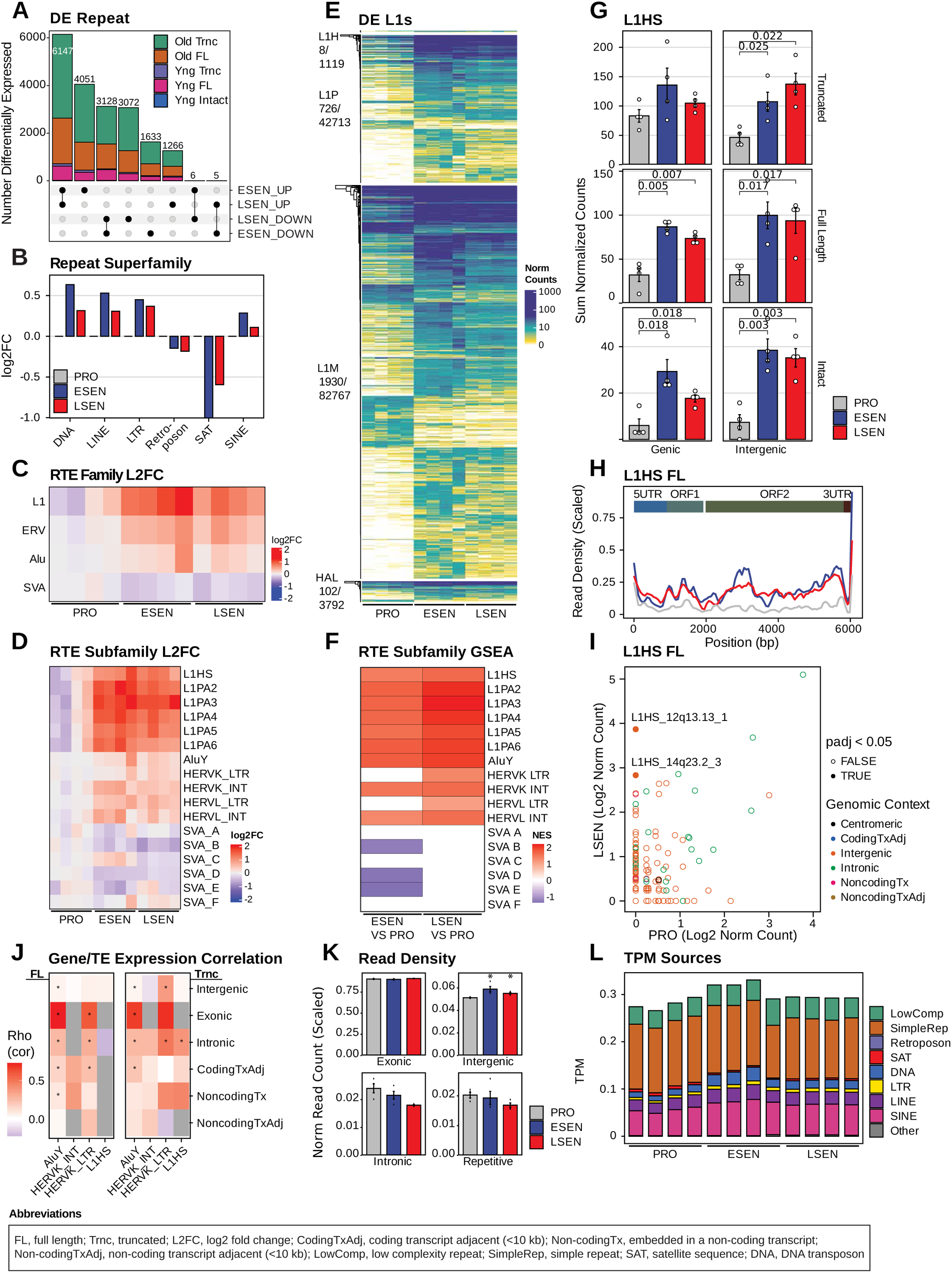
RNA-Seq analysis of TEs. **A** Upset plot of differentially expressed repetitive elements, colored by evolutionary age, element length, and element intactness. **B,C** Log2 fold changes in DESeq2 normalized expression for repetitive element clades: **B** barplot of repeat superfamilies **C** heatmap of RTE families. **D** Heatmap of log2 fold changes in DESeq2 normalized expression for selected RTE subfamilies. **E** Heatmap of DESeq2 normalized expression of differentially expressed (padj <= 0.05) L1 elements. Elements are grouped by clade and hierarchically clustered by expression. Clade labels are followed by a fraction denoting the number of differentially expressed elements over the total number of elements. **F** Gene set enrichment analysis of selected RTE subfamilies for both the ESEN vs PRO and LSEN vs PRO comparisons. Each RTE subfamily is treated as a gene set. **G** Bar plots of total DESeq2 normalized read counts derived from various categories of L1HS elements. **H** Composite expression profile of all full-length L1HS elements. **I** Scatter plot showing log2 DESeq2 normalized counts of every full-length L1HS element, colored by genomic context. **J** Heatmap of Spearman correlation coefficients (Rho) for the log2 fold change in expression of RTEs and neighboring genes. The log2 fold change for each RTE was matched to that of the nearest gene. RTEs are grouped as full-length or truncated, and then by their genomic context relative to the nearest cellular gene. Significant (p<=0.05) associations are denoted by a *. **K** Bar plot of total DESeq2 normalized read counts derived from exonic, intronic, and intergenic compartments, scaled by the total number of counts in each sample**. L** Bar plot showing the distribution of scaled (sum to 1) transcripts per million (TPM) per sample by genomic element class. Unless otherwise stated, statistical significance was assessed by means of a t-test, with panel-level FDR corrected p-values <=0.05*, <=0.01**, and <=0.001***.

Focusing on L1, we observed that many loci spanning all L1 clades were differentially expressed, with the majority being upregulated (Fig. 4E). Most counts were derived from ancient, truncated elements, particularly in the L1M group. This is in keeping with the fact that the majority of L1 elements belong to this group, and reflects the overall loss of heterochromatin in senescence. Of particular interest, however, are the full-length and intact L1HS elements, which retain biological activities, including cDNA generation and retrotransposition. These elements had significantly elevated transcript counts in both ESEN and LSEN (Fig. 4G). Changes in full-length L1HS element expression were not correlated with expression changes in adjacent cellular genes (Fig. 4J), suggesting that they are transcribed from their own promoters. Analysis of full-length L1HS elements at the locus level revealed two significantly upregulated elements, both of which were intergenic, and one of which (14q23.2_3) was intact (Fig. 4I).

Read alignments to full-length L1HS elements spanned the entire element and did not exhibit a 3’ bias, which would indicate the misattribution of truncated element reads to full-length elements (Fig. 4H). Seeing as many TEs reside in introns, differences in intronic read fraction, be this due to differences in intron retention or sequencing library preparation, could lead to spurious conclusions about TE regulation (Faulkner, 2023). We did not observe significant changes in the amount of intronic signal between conditions (Fig. 4K), in fact, intronic signal was decreased in senescent cells. There was, however, a small and significant increase in intergenic signal in both senescent conditions (Fig. 4K). Finally, ESEN showed a small increase in the overall share of the transcriptome derived from repetitive elements (Fig 4L). Taken together, these data document an upregulation of TEs, including young, potentially active RTEs, in ESEN and LSEN conditions.

## Discussion and conclusions

Here we present a pipeline for the analysis of TE RNA-Seq data. Our aim is to render analysis of expression of these elements more tractable to the non-expert. We address known pitfalls in the transcriptomic analysis of these elements by generating an enhanced set of TE annotations, employing up-to-date TE-minded computational methods, and discriminating between signals which originate from autonomous transcripts versus elements transcribed from neighboring genes. To this end, our pipeline makes the following distinctions between elements: genic versus non-genic, evolutionarily young versus old, truncated versus full-length, and intact versus open reading frame (ORF) disrupted. The pipeline can be employed with uniquely mapping reads only, or it can utilize expectation maximization assignments, and individual TE loci can be queried for expression. Furthermore, if provided with Nanopore genome DNA sequences, the pipeline builds a custom reference genome which includes all polymorphic, evolutionarily young elements not included in reference genomes. Taken together, these measures can provide a more accurate assessment of TE expression than is currently possible with standard tools, and should improve our ability to assess whether TEs are contributing meaningfully to a biological problem of interest. As a demonstration, we applied this pipeline to a new dataset from primary human lung fibroblasts and showed that full-length and intact young L1 elements were significantly upregulated in replicative senescence. This is consistent with previous data from our and other groups. Of particular interest was the finding that numerous non-reference L1 elements, which were previously not visible to us, contribute to this upregulation.

### Experimental considerations

Several experimental considerations can help to produce a high quality input dataset for this pipeline. 1) Sequencing read mappability is a function of read-length: a 50bp single-end library will have a much greater number of multi-mapping reads as compared to a 150bp paired-end library [14]. Insofar as TE expression analysis is an important goal, we recommend opting for the longest reads possible. 2) Young L1 subfamilies contain a G-rich stretch of nucleotides in their 3’ UTR which is thought to form a G-quadruplex secondary structure [23]. Polymerases used during sequencing library preparation can struggle with this region, leading to artificially depressed counts. Utilizing reverse transcriptase enzymes with high-processivity tolerant of RNA secondary structures, such as those encoded by group II introns, is therefore preferable. Alternatively, PCR-free approaches such as long-read Nanopore direct RNA sequencing may provide a more accurate assessment of young L1 transcript abundance.

### Software requirements

This pipeline uses the Snakemake [24] workflow manager, and consists of several parts. A main snakefile orchestrates the workflow logic and deploys module level snakefiles which in turn contain the rules that specify each step of the analysis. Conda must be available to create a Snakemake environment. All software dependencies besides Snakemake and Docker/Singularity are packaged into a container which is automatically built and deployed by the pipeline at runtime. Users can alternatively choose to manually build the several conda environments required by the pipeline using the provided yaml environment specifications (this will however be slower and less reproducible). This pipeline was developed on a compute cluster running a RedHat Linux OS and that uses the SLURM workload manager. Nevertheless the use of Docker containers should enable users on other operating systems to run the pipeline without issue.

### Hardware requirements

This pipeline allows for parallel execution of jobs which can occur simultaneously. Consequently, it is highly recommended to execute this pipeline on a compute cluster to take advantage of the parallelization offered by Snakemake. Snakemake is designed to work with many commonly-used cluster workload managers such as SLURM. We provide a default Snakemake profile which is compatible with SLURM, and can be straightforwardly modified for use with other workload managers. Many steps require a substantial amount of RAM (>20 GB) to be available on the system. The Docker container is built to work with both X86 and ARM based CPU architectures.

## Methods

### Overall design

The TE-Seq pipeline consists of two overarching modules, *Annotate Reference (AREF)*, and *Short-read RNA-Seq (SRNA)*. Here we outline the overall workflow and in the next section we provide extended details for selected steps (see Fig. 1 for a schematic outline of TE-Seq).

The AREF module begins by fully annotating a user-provided genome for TE content and identifying functionally important groupings of elements. This enriched annotation set allows the subsequent analysis to probe distinctions in expression between truncated, full-length, and full-length open reading frame intact TEs, as well as to distinguish between intergenic versus intragenic elements. If provided with long-read Nanopore DNA-Seq data derived from the specimens under investigation, this pipeline will call non-reference TE insertions (polymorphic TEs not captured by the reference genome) and create an augmented reference genome containing sequences and associated annotations describing all non-reference TE element insertions. This enables the analysis of polymorphic TE elements, which are likely to include some of the most active elements. For use-case flexibility, the AREF module has a number of workflow modifying parameters. These parameters allow users to choose whether to create new annotations from scratch, to update annotations using long-read sequencing data, or to use existing annotations previously created during a previous run of the pipeline in another project.

Starting with raw sequencing reads, the SRNA module performs standard read-level quality control, genomic alignment, and quantification of gene expression. Repeat-specific tools are then deployed to quantify repeat element expression, either by using only uniquely mapped reads, or, by using the information provided by unique mappings to guide probabilistic assignment of multi-mapping reads to specific TEs. Differential transcript expression of repetitive and non-repetitive elements is assessed using DESeq2. Gene set enrichment analyses are performed for both genes and families of repetitive elements.

### Annotation set creation and refinement

Annotation curation begins with a user-provided reference genome. If a precomputed RepeatMasker.out file is not provided, RepeatMasker (www.repeatmasker.org) will be run on each chromosome to annotate repetitive elements. Next, this RepeatMasker.out file is converted to GTF format with a Bash script, and is then checked for errors and converted to GFF3 and BED file formats. User provided RefSeq gene annotations undergo a similar process, and ultimately RepeatMasker-derived and RefSeq-derived annotations are merged to create a TxDb R object.

Next, RepeatMasker-derived annotations are processed by R scripts to enable downstream analyses. By default, RepeatMasker splits annotations for fragmented repetitive elements. Adhering to ‘tidy’ data conventions, we consolidate these split annotations in an R data frame, taking note of their fragmented status. TEs are then further annotated on the basis of their inclusion in phylogenetic and functional classes. RTEs are grouped at the superfamily, family, and subfamily levels. Elements are denoted as full-length or truncated depending on whether they cover at least 95% of their representative consensus sequence. TE subfamilies are deemed ‘young’ if the average percent divergence (as determined by RepeatMasker) of all subfamily members is less than 3% for LINEs, 8% for SINEs, 12% for ERVs, and 15% for all others. These thresholds are user-tunable parameters, and are set by default, where possible, to denote as ‘young’ those subfamilies capable of retrotransposition (e.g. L1HS and L1PA2 for human LINEs). Evolutionarily young, potentially active elements of two retrotransposon clades, LINEs and ERVs, are further examined below.

Young, potentially active L1 element sequences are analyzed for protein coding potential (in humans: L1HS and L1PA2; in mice: L1Md, L1Tf, L1 Gf and L1A subfamilies; in other species: the least diverged 7 LINE subfamilies). For each full-length sequence, open reading frames (ORFs) are identified. Not all ORFs in all elements will be intact; in fact, many will contain SNPs or indels as a result of genetic drift. To identify consensus ORFs, ORF length frequencies are tabulated, Z-scores are computed, and ORF lengths with a Z-score > 4 are selected for consensus sequence generation. All ORFs are subsequently aligned to these consensus sequences, and their percent amino acid difference is computed. ORFs which are less than 5% diverged are considered intact, and elements are considered intact if they possess a full complement of intact ORFs.

Endogenous retrovirus (ERV) elements are composed of long terminal repeat (LTR) sequences and internal (Int) sequences. RepeatMasker does not annotate whole proviruses, but instead splits them into LTRs and Ints. Most LTRs in genomes are present as ‘solo’ LTRs that do not flank an internal ERV sequence (solo LTRs result from host recombination between two LTRs of a provirus). The TE-Seq pipeline thus distinguishes between proviral and solo LTRs, as well as between 5’LTR-flanked and 5’LTR-deficient ERV sequences. For all LTRs, we determine whether they are flanked by full-length (FL) or truncated (Trnc) Ints (+/− 500bp), thereby generating the following categories: LTR (Solo), 5’LTR (Trnc Int), 3’LTR (Trnc Int), 5’LTR (FL Int), 3’LTR (FL Int). For all Ints, we determine whether they have a 5’LTR and mark them accordingly: Int (Has 5’LTR), or Int (No 5’LTR). Additionally, for young, potentially intact ORF encoding ERV element subfamilies (in the human: HERVK, HERVL, HERVW [25, 26]; in the mouse: MMTV, ETn, IAP, MMVL, MMERGLN, MuRRs, MuLV, MERVL subfamilies [27]; in other species: the least diverged 7 LTR subfamilies), we assign each Int and adjacent (+/− 500bp) LTRs of the same family a unique “proviral_group_id”, such that their counts can be pooled and/or assessed for concordance.

All repetitive elements are examined for their proximity to non-repetitive genetic elements (such as cellular genes). The distance to the most proximal transcript is recorded, and the repetitive element is categorically denoted as either exonic, intronic, coding/non-coding gene proximal (+/− 10 kb from transcript start/end; this threshold is user-tunable), or intergenic.

### Nanopore DNA-Seq

The TE-Seq pipeline allows optional non-reference TE insertion calling if provided with Nanopore DNA-Seq reads. Nanopore basecalling is performed with the Oxford Nanopore Dorado program, and reads are aligned to a reference genome using Minimap2 [28]. Long-read sequencing and alignment quality metrics are compiled using PycoQC [29] and Samtools stats [30]. Aligned DNA reads are then fed to the TLDR program [20], which calls non-reference insertions by examining clusters of clipped-reads whose clipped regions align to a TE consensus sequence. We then filter these insertions to retain only those high-confidence calls which have passed all TLDR filters, and further require that: i) the median MapQ score of supporting reads be 60, ii) the insert has a target-site duplication (TSD) which is a hallmark of retrotransposition, and iii) the insert be supported by at least 5 reads, 2 of which must fully span the insert, and that at least 25% of the reads at the insert site support the insertion. We then append the insert’s consensus sequence (and 30bp of reference flanking region up and downstream of the insert) to the provided reference genome where it becomes a standalone, mappable contig. RepeatMasker then annotates these appended contigs for repetitive element content, and these non-reference insertion annotations are merged with the reference TE RepeatMasker set for further processing as described in the previous section. If provided with a species-specific database of TE polymorphisms, TLDR annotates whether insertions are ‘known’ or ‘novel’. TLDR provides such databases for human and mouse (see [20], Table S4 for more information on the 17 datasets comprising this database). Additionally, we optionally call SNPs using the PEPPER pipeline [31], and incorporate high-quality SNP calls into the reference genome allowing for more accurate mapping of reads overlapping these polymorphisms.

If Nanopore DNA sequences are not available, the problem of non-reference TEs can nevertheless be attenuated by using the most up-to-date reference genomes, such as telomere-to-telomere assemblies [21] which provide a more complete map of TE insertions, especially in hard to assemble regions such as centromeres.

### RNA-Seq analysis

Sequencing reads are trimmed using fastp [32] and aligned to the (*custom*) reference genome reference with STAR [33], allowing for up to 100 multiple mapping locations per read. This allowance for multimappers aids in accurately assigning TE-aligned reads to the correct TE locus. Gene counts are produced using featureCounts [34] and the user-provided RefSeq transcript annotation (in the present case, the October 2023 revision, GCF_009914755.1). Repetitive element count estimates are produced with the Telescope tool [18] and the custom RepeatMasker derived annotation resulting from the AREF module. This tool allows for multi-mapping reads overlapping TEs to be assigned to the best supported locus via an iterative expectation maximization (EM) algorithm (see Telescope methods). We obtain two sets of TE counts: one conservative estimate of locus expression which allows only for uniquely-mapped reads to contribute to total counts, and one more relaxed (and potentially accurate, as claimed in [18]) set of counts which incorporates multi-mapping reads. This inclusion of multi-mapping reads is particularly important for young TE elements, which are highly similar in sequence and contain long stretches of non-uniquely mappable sequences. While overall a less biased strategy than only considering uniquely mapping reads [18], EM re-assignment of multi-mapping reads is imperfect, and element-level read estimates should be interpreted as best guesses when they are comprised of a substantial fraction of multi-mapping reads.

Count normalization for genes and repetitive elements is performed using DESeq2’s ‘median of ratios’ method [35]. Size factors are estimated by setting the ‘controlGenes’ parameter of the ‘estimateSizeFactors’ factors to include only RefSeq genes. This is done so as to prevent the very large number of lowly-expressed repetitive elements in a RepeatMasker-derived annotation from adversely biasing median-based count normalization. Differential expression is then assessed using DESeq2’s negative binomial linear modeling. Batch effect correction is implemented by introducing a batch variable in the sample specification file. If sequencing data contain batch effects, one must be mindful that batch effect correction is an imperfect procedure which can in itself introduce a bias [36, 37]. This problem is particularly acute when biological conditions of interest are imbalanced with respect to batch. The TE-Seq pipeline accounts for batch effects in two places: the first is in DESeq2’s linear model, and the second is during the production of batch-corrected count tables using limma [38]. DESeq2’s linear modeling will better handle the batch effect, and its p-value estimates (for individual genes and repetitive elements) will be more reliable than any statistics produced downstream of limma ‘batch-corrected’ counts (comparisons between repetitive element family total counts). Therefore, in the context of an unbalanced dataset, TE family comparisons based on adjusted counts need to be interpreted with care, and in the light of non-batch adjusted results. For a more thorough discussion of the issue, readers may consider [37].

Normalized counts and differential expression results are next analyzed with a set of R scripts. Example products of this analysis are shown in the results section. Analyses at both the individual gene/TE level, as well as at the gene set/TE family level are performed. TE groupings are assessed for enrichment using both gene set enrichment analysis (GSEA) and pooled-count strategies. GSEA can detect small, concerted changes in expression across a family of related TEs. Normalized TE count pooling at the family or functional level (and subsequent assessment for differential expression by means of a t-test) allows for a TE family to be counted as significantly overexpressed if only a small number of its members are dramatically overexpressed.

Correlations in expression between adjacent gene-TE pairs are established to shed light on potential bidirectional gene-regulatory influences. Read-mapping to select TE families is also examined by means of composite bigwig tracks, which show read density across a consensus element’s length, revealing potential biases in alignment to particular TE sequence regions/features.

Finally, the pipeline produces an html report consolidating the most insightful visualizations. Plots are stored as PDFs such that they can be resized and modified as needed in vector graphics software such as Adobe Illustrator.

## Acknowledgements

The authors would like to thank Jess Anderson for lab management and technical assistance, and members of the Sedivy lab and the Center on the Biology of Aging for feedback and support. We are grateful to the Brown Center for Computation and Visualization (CCV) team for their management of the OSCAR high performance computing cluster which was used throughout this work.

## Authors’ contributions

M.M.K. developed all code and figures, and performed long-read Nanopore DNA sequencing of LF1 cells. R.L.K. provided short-read RNA sequencing of LF1 cells. J.M.S. supervised the project. M.M.K. and J.M.S coauthored the manuscript.

## Availability of data and materials

Instructions for pipeline installation, configuration, and deployment are documented at the project GitHub page: https://github.com/maxfieldk/TE-Seq. This page also provides instructions for the acquisition of human and mouse reference genome annotations. A Zenodo archive of the software can be found at: <u>10.5281/zenodo.14680966</u>. RNA-Seq data and Nanopore DNA-Seq data have been deposited in the Gene Expression Omnibus (GEO) with accession numbers GSE280365 and GSE268488, respectively. Down-sampled versions of these data are made available for pipeline testing purposes and can be found at the following GitHub page: https://github.com/maxfieldk/TE-Seq_test_rawdata. Any other data or information relevant to this study are available from the corresponding author upon reasonable request.

## Declarations

### Ethics approval and consent to participate

Not applicable

### Consent for publication

Not applicable

## References

1. Almojil D, Bourgeois Y, Falis M, Hariyani I, Wilcox J, Boissinot S. The Structural, Functional and Evolutionary Impact of Transposable Elements in Eukaryotes. Genes. 2021; 12:918.

2. Baduel P, Quadrana L. Jumpstarting evolution: How transposition can facilitate adaptation to rapid environmental changes. Curr Opin Plant Biol. 2021; 61:102043.

3. Rodriguez-Martin B, Alvarez EG, Baez-Ortega A, Zamora J, Supek F, Demeulemeester J, Santamarina M, Ju YS, Temes J, Garcia-Souto D, et al. Pan-cancer analysis of whole genomes identifies driver rearrangements promoted by LINE-1 retrotransposition. Nat Genet. 2020; 52:306–319.

4. Kofler R, Nolte V, Schloetterer C. Tempo and Mode of Transposable Element Activity in Drosophila. PLoS Genet. 2015; 11:e1005406.

5. Konkel MK, Walker JA, Batzer MA. LINEs and SINEs of Primate Evolution. Evol Anthropol. 2010; 19:236–249.

6. Bourque G, Burns KH, Gehring M, Gorbunova V, Seluanov A, Hammell M, Imbeault M, Izsvák Z, Levin HL, Macfarlan TS, et al. Ten things you should know about transposable elements. Genome Biology. 2018; 19:199.

7. Frost B, Dubnau J. The Role of Retrotransposons and Endogenous Retroviruses in Age-Dependent Neurodegenerative Disorders. Annu Rev Neurosci. 2024; 47:123–143.

8. Gorbunova V, Seluanov A, Mita P, McKerrow W, Fenyo D, Boeke JD, Linker SB, Gage FH, Kreiling JA, Petrashen AP, et al. The role of retrotransposable elements in ageing and age-associated diseases. Nature. 2021; 596:43–53.

9. Mendez-Dorantes C, Burns KH. LINE-1 retrotransposition and its deregulation in cancers: implications for therapeutic opportunities. Genes Dev. 2023; 37:948–967.

10. Konkel MK, Batzer MA. A mobile threat to genome stability: The impact of non-LTR retrotransposons upon the human genome. Semin Cancer Biol. 2010; 20:211–221.

11. Burns KH, Boeke JD. Human transposon tectonics. Cell. 2012; 149:740–752.

12. Lanciano S, Cristofari G. Measuring and interpreting transposable element expression. Nat Rev Genet. 2020; 21:721–736.

13. Kaul T, Morales ME, Sartor AO, Belancio VP, Deininger P. Comparative analysis on the expression of L1 loci using various RNA-Seq preparations. Mob DNA. 2020; 11:2.

14. Teissandier A, Servant N, Barillot E, Bourc’his D. Tools and best practices for retrotransposon analysis using high-throughput sequencing data. Mobile DNA. 2019; 10:52.

15. Lerat E, Fablet M, Modolo L, Lopez-Maestre H, Vieira C. TEtools facilitates big data expression analysis of transposable elements and reveals an antagonism between their activity and that of piRNA genes. Nucleic Acids Res. 2017; 45:e17.

16. Criscione SW, Zhang Y, Thompson W, Sedivy JM, Neretti N. Transcriptional landscape of repetitive elements in normal and cancer human cells. BMC Genomics. 2014; 15:583.

17. McKerrow W, Fenyö D. L1EM: a tool for accurate locus specific LINE-1 RNA quantification. Bioinformatics. 2020; 36:1167–1173.

18. Bendall ML, Mulder Md, Iñiguez LP, Lecanda-Sánchez A, Pérez-Losada M, Ostrowski MA, Jones RB, Mulder LCF, Reyes-Terán G, Crandall KA, et al. Telescope: Characterization of the retrotranscriptome by accurate estimation of transposable element expression. PLOS Comp Biol. 2019; 15:e1006453.

19. Brown JP, Wei W, Sedivy JM. Bypass of senescence after disruption of p21CIP1/WAF1 gene in normal diploid human fibroblasts. Science. 1997; 277:831–834.

20. Ewing AD, Smits N, Sanchez-Luque FJ, Faivre J, Brennan PM, Richardson SR, Cheetham SW, Faulkner GJ. Nanopore Sequencing Enables Comprehensive Transposable Element Epigenomic Profiling. Molecular Cell. 2020; 80:915–928.e915.

21. Nurk S, Koren S, Rhie A, Rautiainen M, Bzikadze AV, Mikheenko A, Vollger MR, Altemose N, Uralsky L, Gershman A, et al. The complete sequence of a human genome. Science. 2022; 376:44–53.

22. Hernandez-Segura A, Nehme J, Demaria M. Hallmarks of Cellular Senescence. Trends Cell Biol. 2018; 28:436–453.

23. Sahakyan AB, Murat P, Mayer C, Balasubramanian S. G-quadruplex structures within the 3’ UTR of LINE-1 elements stimulate retrotransposition. Nat Struct Mol Biol. 2017; 24:243–247.

24. Mölder F, Jablonski KP, Letcher B, Hall MB, Tomkins-Tinch CH, Sochat V, Forster J, Lee S, Twardziok SO, Kanitz A, et al: Sustainable data analysis with Snakemake. F1000Research; 2021.

25. Garcia-Montojo M, Doucet-O’Hare T, Henderson L, Nath A. Human endogenous retrovirus-K (HML-2): a comprehensive review. Crit Rev Microbiol. 2018; 44:715–738.

26. Jakobsson J, Vincendeau M. SnapShot: Human endogenous retroviruses. Cell. 2022; 185:400–400.e401.

27. Stocking C, Kozak CA. Endogenous retroviruses. Cellular and Molecular Life Sciences: CMLS. 2008; 65:3383–3398.

28. Li H. Minimap2: pairwise alignment for nucleotide sequences. Bioinformatics. 2018; 34:3094–3100.

29. Leger A, Leonardi T. pycoQC, interactive quality control for Oxford Nanopore Sequencing. Journal of Open Source Software. 2019; 4:1236.

30. Danecek P, Bonfield JK, Liddle J, Marshall J, Ohan V, Pollard MO, Whitwham A, Keane T, McCarthy SA, Davies RM, Li H. Twelve years of SAMtools and BCFtools. GigaScience. 2021; 10:giab008.

31. Shafin K, Pesout T, Chang PC, Nattestad M, Kolesnikov A, Goel S, Baid G, Kolmogorov M, Eizenga JM, Miga KH, et al. Haplotype-aware variant calling with PEPPER-Margin-DeepVariant enables high accuracy in nanopore long-reads. Nat Meth. 2021; 18:1322–1332.

32. Chen S, Zhou Y, Chen Y, Gu J. fastp: an ultra-fast all-in-one FASTQ preprocessor. Bioinformatics. 2018; 34:i884–i890.

33. Dobin A, Davis CA, Schlesinger F, Drenkow J, Zaleski C, Jha S, Batut P, Chaisson M, Gingeras TR. STAR: ultrafast universal RNA-seq aligner. Bioinformatics. 2013; 29:15–21.

34. Liao Y, Smyth GK, Shi W. featureCounts: an efficient general purpose program for assigning sequence reads to genomic features. Bioinformatics. 2014; 30:923–930.

35. Love MI, Huber W, Anders S. Moderated estimation of fold change and dispersion for RNA-seq data with DESeq2. Genome Biol. 2014; 15:550.

36. Leek JT, Scharpf RB, Bravo HC, Simcha D, Langmead B, Johnson WE, Geman D, Baggerly K, Irizarry RA. Tackling the widespread and critical impact of batch effects in high-throughput data. Nat Rev Genet. 2010; 11:733–739.

37. Nygaard V, Rødland EA, Hovig E. Methods that remove batch effects while retaining group differences may lead to exaggerated confidence in downstream analyses. Biostatistics. 2016; 17:29–39.

38. Ritchie ME, Phipson B, Wu D, Hu Y, Law CW, Shi W, Smyth GK. limma powers differential expression analyses for RNA-sequencing and microarray studies. Nucleic Acids Res. 2015; 43:e47.

